# Anti-fibrotic Effects of Atorvastatin Delivered via Keratin-based Hydrogels

**DOI:** 10.1101/2024.12.09.627550

**Authors:** Evan Carroll, Andrew Tarabokija, Henna Chaudhry, Allison Meer, Roche C. de Guzman

**Affiliations:** Bioengineering Program, Department of Engineering, Hofstra University, Hempstead, NY 11549; Department of Chemistry, Hofstra University, Hempstead, NY 11549; Department of Biology, Hofstra University, Hempstead, NY 11549

## Abstract

Implantations of degradable biomaterials for drug delivery or restoration and regenerative medicine cause fibrosis and acute inflammation which may lead to chronic side effects. Additionally, the fibrous encapsulation can itself inhibit the local cells from proliferating and forming new healthy tissue. Current drug delivery models require a new method to decrease the adverse effects following implantation of biomaterials in patients and inhibit the activation signaling sent to quiescent fibroblasts. This research details the novel chemical and physical properties of reduced keratin (KTN) hydrogels extracted from residual human hair with sheared cuticle layers, obtained from barbershops and beauty salons. KTN hydrogels with and without calcium ions conjugated with atorvastatin (Ator), which is thought to affect TGF-ß signaling pathways directly by competitively inhibiting key signal transmitters and cellular responses, were investigated via electron microscopy, absorbance spectroscopy, and computational modeling and simulation to observe the proliferative inhibition of fibroblastic cells in comparison to the alginate hydrogel controls. Hydrogel extracts were tested for non-cytotoxic effects via L929 adipose fibroblast cell culture following ISO 10993-5 standard for safety of medical device biomaterials. Two cell models were exposed to increasing serial concentrations of Ator to evaluate the half maximal effective concentration (EC_50_). For quiescent fibroblasts, Ator’s EC_50_ was observed at 368 μM in PBS. For mesenchymal stem cells (MSCs), Ator’s EC_50_ was observed at 209 μM in PBS. The mass of drug was normalized in all groups after calculating their timed absorption and release, and KTN hydrogels conjugated with calcium ions and Ator (ATCK) completely absorbed the drug < 0.5 h while KTN hydrogels with Ator alone (ATKR) completely absorbed the drug at 1h. After 7d in PBS, ATCK released Ator at 2.2 ± 0.7% while Ator in calcium alginate spheres (ATLG) was released at 7.8 ± 1.1% and ATKR released 11.2 ± 4.3% of drug. On-going research focuses on other active fibroblastic cell types and *in vivo* experimentation to observe foreign body response in subcutaneous mouse tissue.

## 1. Introduction

The development of advanced drug delivery systems has revolutionized the field of medicine and offers targeted, controlled, and sustained release of therapeutic agents. Natural and synthetic hydrogels are considered one of the leading materials for drug delivery carriers, and naturally obtained keratin-based hydrogels derived from human hair represent a cutting-edge discovery in the field of functional biomaterials. Keratin (KTN) and keratin associated proteins (KAP), found abundantly in human hair and nails, are renewable, degradable biopolymers whose byproducts are non-toxic and account for > 95% of the mass of each human hair fiber. Each fiber consists of a thin, scaly cuticle layer that protects the cortex, constituting the bulk of hair proteins^1,2^. Chemical reduction processes are widely used to expose cortical matrices and modify their molecular scaffolds for gel formation^1,3^. Our research group and others^3-9^ have determined the effectiveness of soluble and extractable protein fractions of KTN and KAP in animal models for tissue engineering and personalized medicine, but this study focuses on the insoluble fractions, initially deemed as “waste” products, that can enhance wound healing and controlled drug transport for topical and localized delivery.

Statins are commonly prescribed medications used to decrease cholesterol by inhibiting 3-hydroxy-3-methyl-glutaryl-coenzyme A reductase. Importantly, they have pleiotropic anti-fibrotic effects, possibly through the transforming growth factor beta (TGF-ß) signaling pathway which either prevents the activation of quiescent fibroblast or transdifferentiation of progenitors into myofibroblast^10-14^. The anti-inflammatory effects enhance the drug-conjugated hydrogels to create a conducive environment for tissue repair while also promoting angiogenesis to improve wound healing. The bioactivities of Atorvastatin (Ator, *Atorvastatin calcium tablets USP, Dr. Reddy’s Laboratories Limited*), were investigated when delivered locally via KTN hydrogels as novel drug transport vehicles for disease management. This localized action reduces systemic side effects and targets Ator’s therapeutic effects more effectively at the injury site. **The goal is to control drug loading and release of Ator to produce a therapeutic effect for scar tissue inhibition**.

KTN hydrogels can not only sequester Ator but also crosslink with other drugs and biologics such as growth factors, antibiotics, or anti-inflammatory drugs and play oncogenic roles^7,15-18^. This opens the door for more multi-functional drug carriers and offers a broader range of therapeutic benefits, such as antibacterial defense with tissue regeneration and inflammation control. The properties of Ator-conjugated KTN hydrogels can be tailored by altering the Ator release rate or the hydrogel stiffness to more effectively provide personalized medicine and address patient-specific needs, including the severity of the wound, skin type, or underlying conditions.

## 2. Materials and Methods

### 2.1 Preparation and Characterization of Keratin Hydrogels

Human hair samples were obtained from local barbershops and hair salons (*East Meadow, NY*). The hair was washed with soap to remove grime, dehydrated in ethanol (EtOH), and delipidized in 2:1 volumes (V) of chloroform to methanol at 37°C, overnight (O/N) in an environmental shaker at 140 rpm to remove their outer fat layers on the cuticle. To oxidize the melanin in hair, a bleaching mix of 1 V bleach powder (*Perfect Blond, Blond Forte, Medley, FL*) to 2 V developer (*Crème 40 V, Clairol Professional, Petit-Lancy, Geneva, Switzerland*) was lathered on hair spread and wrapped in aluminum foil for 40 minutes and rinsed with water, repeated twice. The oxidized, delipidized hair was separated into batches that were incubated at 50 mg/mL in a thioglycolic acid reducing solution (0.5M TGA, pH 11 adjusted) shaking at 37°C for 24 hours to break the disulfide bonds and release small proteins. The insoluble fraction was sieved and transferred into a buffer solution (0.1M tris) and placed for two hours in the environmental shaker, accounting for the new total volume. The insoluble hair was again sieved and transferred into deionized water for two hours in the environmental shaker to wash away any leftover chemicals. This process was repeated once more to reduce the insoluble hair content. The insoluble hair contents remaining were washed using dialysis tubing to remove any unwanted chemicals and swell the contents to become more gel-like. Dialysis continued for three weeks, and the water content was changed every three days.

To observe the effects of the reduction process and visualize the microstructure and pores of the hydrogels scanning electron microscopy (SEM) was taken. Samples were dehydrated in increasing EtOH (50%, 70%, 95%, and 100% for 5 minutes each). After sputter-coating (*EMS-550 Sputter Coater, Electron Microscopy Sciences, Hatfield, PA*) their surfaces with gold for electron conductivity, KTN hydrogels were imaged via a Q250 SEM (*FEI, Hillsboro, OR*) at 30 kV. Since keratin proteins make up most proteins in human hair, the hydrogel porosity was quantified by removing excess liquid and comparing the mass of the gels (n = 4) before and after total drying at 100°C in a convection drying oven. The hydrogel swell ratio was calculated by measuring the difference in mass between hydrogels (n = 4) equilibrated in PBS and then submerged in deionized water. Hydrogel surface degradability was quantified by taking the initial gel mass (n = 4) before adding 400 μL of PBS to each group and taking their final masses after removing the 400 μL PBS at days 0, 0.25, 1, 3, and 7.

### 2.2 Experimental and Control Hydrogelation

To increase Ator’s gel retention, positively charged calcium ions were mixed to electrostatically-conjugate the negatively charged KTN and Ator. Freshly dialyzed KTN gels from insoluble hair solutions infused with (CKRT) and without calcium (KRT) were prepared. CKRT gels were submerged in 400 mM calcium chloride solutions for a week to absorb the maximum amount of calcium ions. Calcium alginate spheres served as the control hydrogels since they are widely used biomaterials in drug delivery applications. They were synthesized by slowly dripping 2% sodium alginate droplets into 400 mM calcium chloride. All experimental hydrogels included 200 μg/mL or 368 μM Ator in PBS in their matrices.

### 2.3 Drug Quantification and In Vitro Pharmacology

Spectrophotometry was used to measure the concentration of Ator absorbed and released using a nano UV-Vis spectrophotometer (*NanoDrop™ 2000, Thermo Fisher Scientific, Waltham, MA*). Its description and pharmacokinetics are tabulated in **Table 1**. Increasing Ator concentrations were detected via spectral scanning over the range of visible wavelengths. The hydrogels and calcium solutions were tested to assure no interference at the optimal wavelength.

**Table 1.**
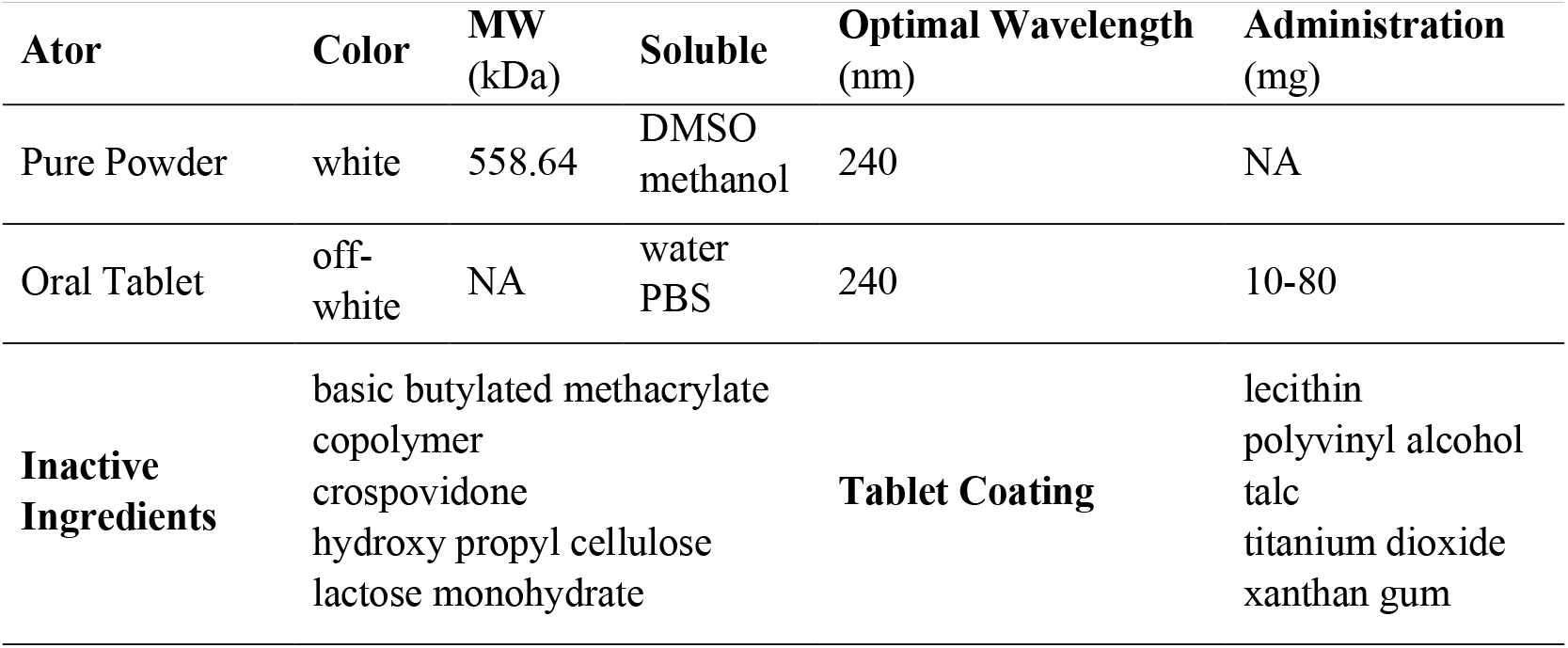
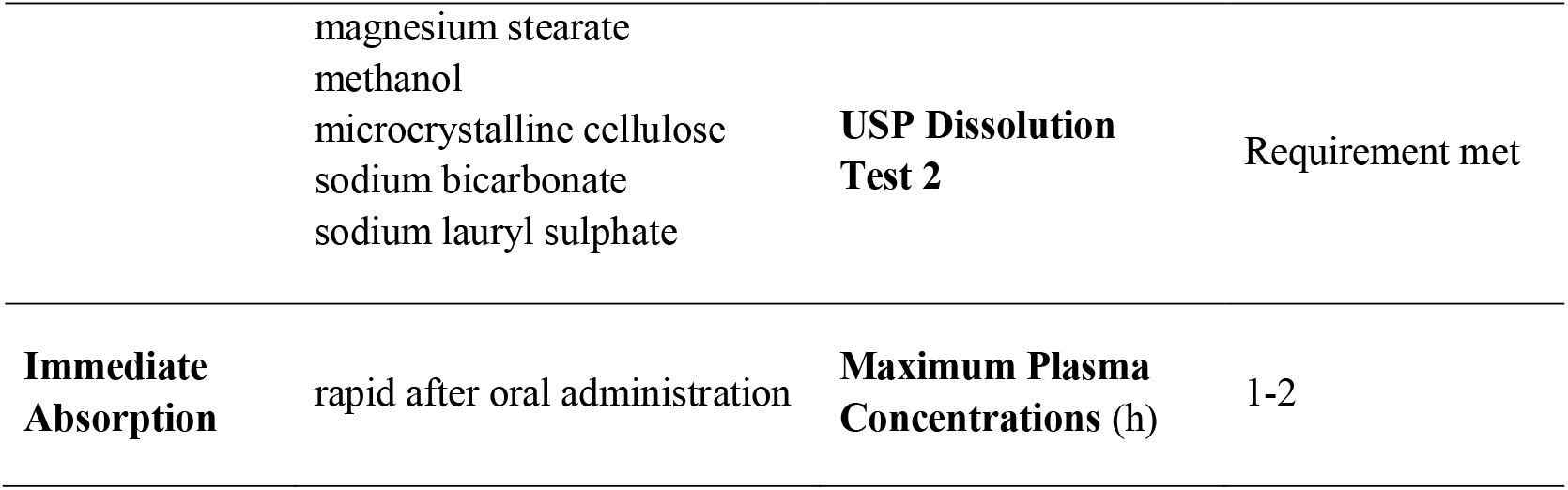
Drug description and Pharmacokinetics of Atorvastatin calcium from the Dr. Reddy’s Laboratories Limited Safety Data Sheet^19^. MW = molecular weight. NA = not applicable.

The diffusion of Ator through the KTN hydrogel matrix is influenced by the pore size which dictates the release process. When the pore size exceeds the Ator’s radius (r_gel_/r_Ator_ > 1), the drug release is governed by diffusion. The diffusivity, *D*, depends on the radius of Ator (r_Ator_) and the viscosity of the solution (η) via the Stokes–Einstein equation:

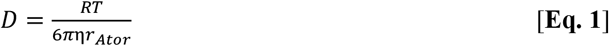

where *R* is the gas constant and *T* is the absolute temperature. The radius of the drug molecule correlates with its molecular weight, meaning macromolecular drugs have larger radii and experience slower diffusion^20^. KRT and CKRT gels were soaked in a 200 μg/mL Ator solution in PBS for one week. The absorbance of Ator in the solution was measured at nine time points (0, 0.25, 0.5, 1, 2, 4, 24, 96, 168 h) to quantify the drug diffusion into the gel and its cumulative mass ratio. The Ator-diffused gels were then placed in PBS to observe the release profiles at eight time points (0, 0.25, 0.5, 1, 2, 24, 96, 168 h). Measured data was then simulated in COMSOL Multiphysics® (*COMSOL AB, Stockholm, Sweden*) to observe how the molecular size, solubility, and keratin crosslinking affected the Ator diffusion coefficient and the rate of absorption:

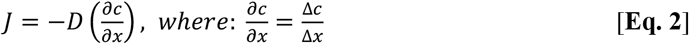

where 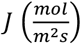 is the diffusion flux, 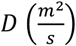 is the Ator diffusion coefficient in the hydrogel, and 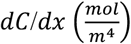 is the Ator concentration gradient through the vertical distance in the hydrogel. To estimate the cumulative mass absorption rate, 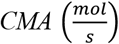, the diffusion flux is multiplied by the cross-sectional area, *A* (*m*^2^), of the hydrogel that the drug is diffusing:

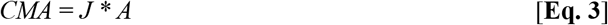

### 2.4 In Vitro Cell Line Culture

Ator’s dose-response effect to fibroblast precursors were evaluated using mesenchymal stem cells’ (MSCs, ASC52telo, ATCC) viability. This *in vitro* dose-response biocompatibility test was also employed on L-929 mouse adipose areolar subcutaneous fibroblasts line (NCTC clone 929, ATCC, Manassas, VA) according to ISO 10993-5: Biological evaluation of medical devices, Tests for *in vitro* cytotoxicity, Annex C: MTT assay. The cytotoxicity of all hydrogel groups was also verified through extract tests on L-929 fibroblasts. Both MSC and fibroblast cell lines were cultured in Dulbecco’s Modified Eagle’s Medium (DMEM) with 5% (V/V) and 10% (V/V) fetal bovine serum, respectively, and 1× antibiotic-antimycotic agent in a 96-well plate at 100-mL with 10^4^ cells/well (n = 3) for 24 h in a mammalian cell incubator (37 °C, 5% CO2, >90% relative humidity). L-929 and MSC inhibition was observed at increasing Ator concentrations, *c*, (0, 3.125, 6.25, 12.5, 25, 50, 75, 100, 125, 150, 175, 200, 225 ug/mL) in DMEM. The dose response, *E*, of the negative sigmoidal inhibition curve was calculated using the parameters: *c, E*_*max*_, the maximal possible effect, *EC*_*50*_, the half maximal effective concentration, and *n*, Hill’s coefficient which is negative for cell toxicity, and plotted using Hill’s model:

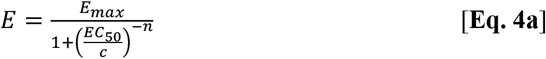

The parameters were found to produce the best fit by minimizing the sum of square errors between the predicted values from the Hill equation and the measured data points in MATLAB (*The MathWorks Inc*., *Natick, MA*).

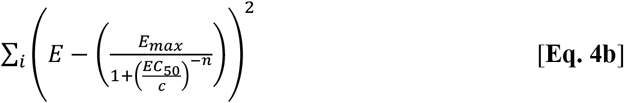

Hydrogels were shaken with 50% EtOH for 1 minute and immediately submerged in sterile water. After removing the water after 5 minutes, the gels were soaked in DMEM at 3 cm^2^/mL material surface area to DMEM volume ratio and incubated for 3 days at 37°C with shaking for extraction of any leachable substances, in accordance to ISO 10993-12: Sample preparation and reference materials, including latex (and 0.6% dimethyl sulfoxide added to cells) as the positive (+) and polypropylene as the negative (–) biomaterial cytotoxicity controls. 100-mL of these extracts at different serial dilutions (1, 0.5, 0.25, 0.125, and 0 × blank in DMEM) were aspirated on to cultured cells for 20 hours. A non-hydrogel control was 200 μg/mL Ator aspirated on to cells to observe how the cells reacted to quick delivery. The cells were incubated for an additional 4 hours after aspirating 50 mL of 1 mg/mL (3-(4,5-dimethylthiazol-2-yl)-2,5-diphenyltetrazolium bromide) (MTT) in DMEM, and the culture medium was removed to add isopropanol to dissolve the formazan crystals and compare the absorbance between experimental and control groups at 570-nm.

### 2.5 Statistical Methods

Images were processed in ImageJ and PowerPoint (Microsoft). Data and graphs were evaluated and generated in Excel. Technical and biological replicate values were averaged and reported as means (AVE) ± 1 standard deviations (STD). Student’s t-test and analysis of variance (ANOVA) with Tukey-Kramer were used for multiple comparison using a probability (p) < 0.05 deemed significantly different.

## 3. Results

### 3.1 Keratin Hydrogel Properties

The processed residual hairs were viscous and relatively dense in consistency after dialysis and exhibited a pale-yellow color (**Fig. 1A**) with soft strands on which the Ator macromolecules would crosslink. The calcium alginate gels were clear and spherical (**Fig. 1B**), and both gels were easily compressible and stretchable. The final insoluble hair mass yield from the initial mass of KTN was 56%. KTN hydrogels exhibited a mass degradation rate of 2.07% per day (**Fig. 1C**), and after linear extrapolation under exaggerated conditions, they will exhibit complete degradation relative to 0% at a two-month timepoint. Once the hydrogels were completely dry (**Fig. 1D**), their density of 1.022 g/mL was experimentally calculated from a protein percent yield of 10.97 ± 0.91%. The porosity is the space that water took up in the hydrogel, 89.03 ± 0.91%, which was indirectly calculated. The gel swell ratio was calculated to be 108.10 ± 9.48% greater in water than in PBS. Hydrogel topography and cross-sectional area confirmed the efficacy of the hair reduction process. The KTN cuticle (**Fig. 2A**), now wrinkled and irregular due to loss of cortical contents (**Fig. 2B**), remained present through most of the hair matrix, but had large gaps (**Fig. 2C**) torn down the length of hair strands to reveal loose cortical matrices (**Fig. 2D**). Electron micrographs analyzed in ImageJ exhibited a void space area of 55.7% and each hair with a diameter of 82.5 ± 44.5 μm.

**Fig 1.**
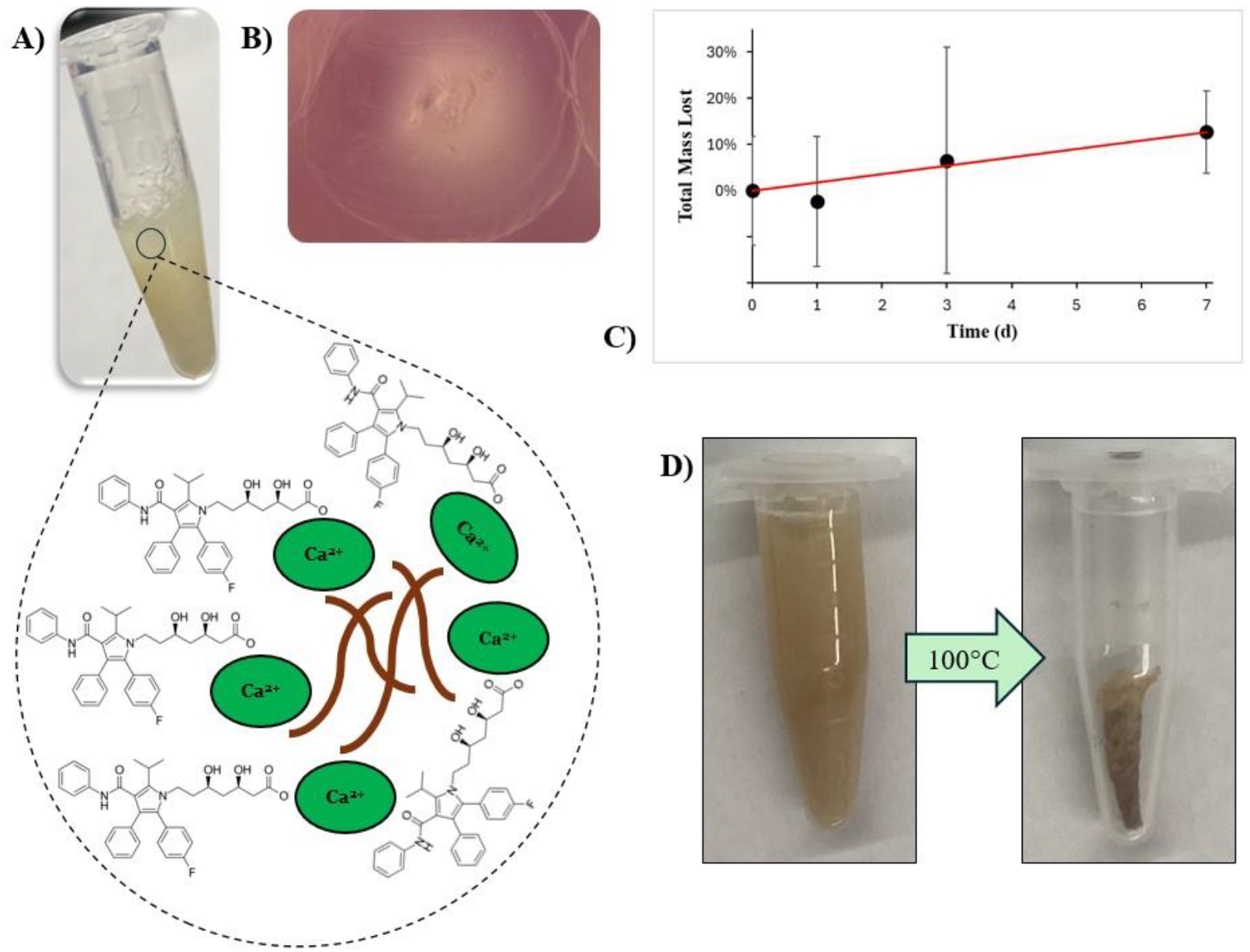
**A)** Processed residual hair hydrogel (KRT, CKRT, ATKR, & ATCK groups look identical). ATKR & ATCK hydrogels have KTN matrices that sequester Ator and calcium ions (ATCK > Ca > ATKR). **B)** Microscopy image of an alginate sphere (CALG & ATLG groups look identical, d = 3 mm). **C)** KTN degradation rate (r = 2.07% per day); 100% degradation projected after 49 days. **D)** KTN hydrogel at swollen equilibrium state (left) then dried completely in a convection oven at 100°C (right) for protein yield.

**Fig 2.**
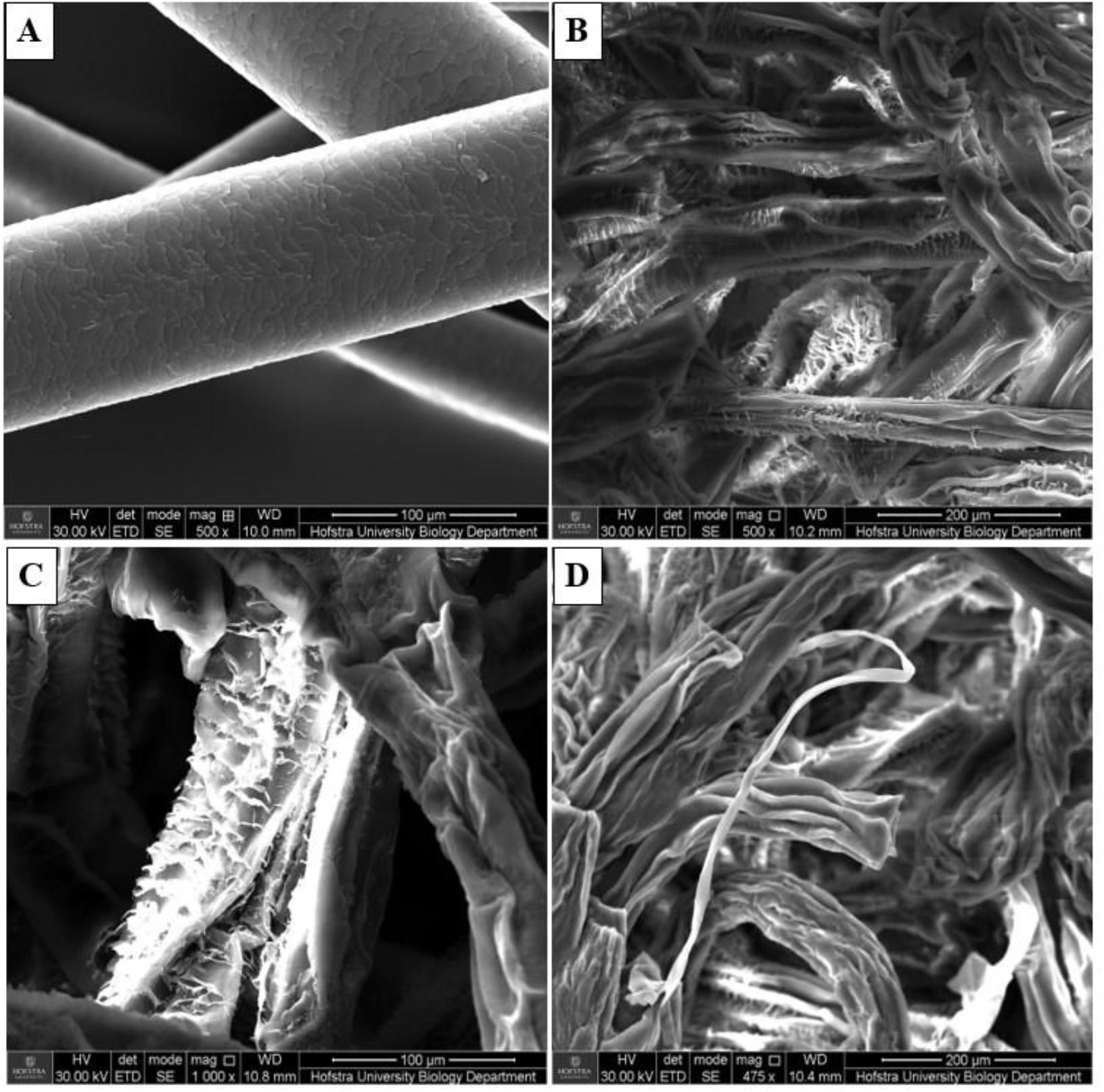
SEM imaging of **A)** untreated regular human hair with the cuticle intact, **B)** treated KTN hydrogel matrix with sheared cuticles and exposed inner cortical bulk, **C)** the shriveled structure of processed residual hair with cuticle fragments, and **D)** potential macrofibril strands within the treated KTN hydrogel matrix which are beneficial for biomaterial scaffolds and drug sequestration.

The keratin hydrogel properties are summarized in **Table 2**.

**Table 2.** Physical properties of processed human hair to display its utility in forming drug-conjugated hydrogels.

| Processed residual hair | Color and Appearance | SEM Topography | Void Space | Fiber Diameter ($\mu m$) |
| --- | --- | --- | --- | --- |
| 56% | pale-yellow, wavy | wrinkled, sheared cuticle, exposed cortex | 55.7% | $82.5 \pm 44.5$ |

| Density<br>(g/mL) | Protein Yield | Swell Ratio<br>(PBS → H<sub>2</sub>O) | Porosity | Mass<br>Degrada-<br>tion Rate |
| --- | --- | --- | --- | --- |
| 1.022 | 10.97 ± 0.91% | 108.10 ± 9.48% | 10.97 ±<br>0.91% | 2.07% |

### 3.2 In Vitro Drug Pharmacokinetics

Ator was absorbed into KRT, CKRT (**Fig. 3A, top**), and CALG hydrogel groups (ATKR, ATCK, and ATLG, respectively) to observe *in vitro* pharmacological properties such as the mass diffusivity and slow-release effects, and the optimal wavelength to measure Ator concentrations is 240-nm. The gels potential for injectability was also observed before and after Ator sequestration (**Fig. 3A, bottom**). ATKR fully absorbed 1.8 μg/mg Ator after 1 h while ATCK absorbed 1.8 μg/mg Ator after 15 min (**Fig. 3B**). At 15 min, Ator had absorbed twice as fast into ATCK as ATKR while Ator was immediately sequestered into ATLG at the time of synthesis. This experimental data was used to simulate the drug-hydrogel pharmacokinetics and to calculate cumulative mass absorption (CMA). The cumulative mass release (CMR) profile at 24 h exhibited a 5-fold increase of Ator release between ATCK and ATKR and a 6.25-fold increase of Ator between ATCK and ATLG (**Fig. 3C**). Meanwhile, after 7 d the CMR profile shows a greater release of ATKR (5-fold) than ATLG (3.8-fold) when compared to ATCK.

**Fig 3.**
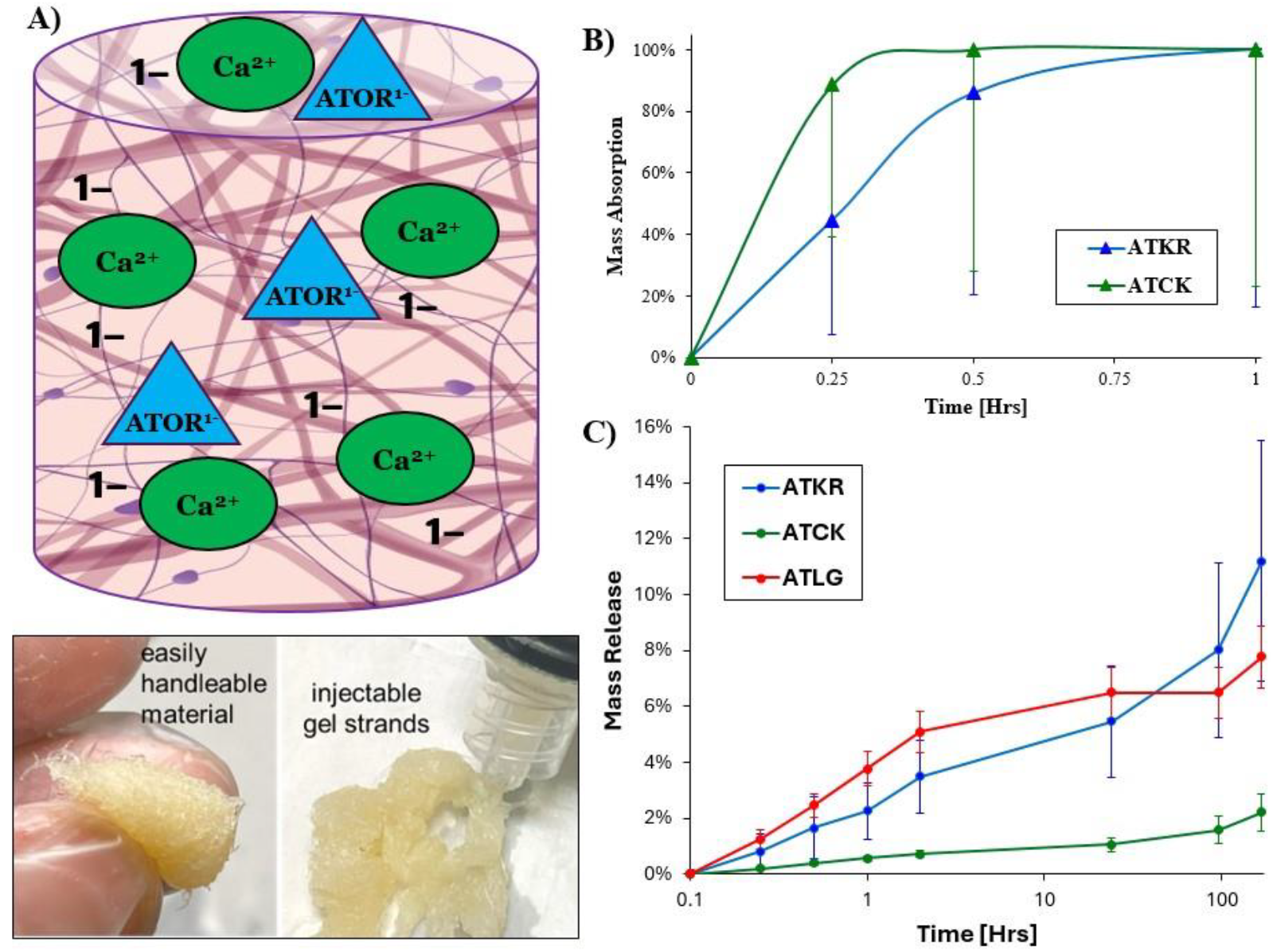
**A)** ATKR or ATCK hydrogel makeup with electrostatic-conjugation (top) and structural integrity while exhibiting easy handleability and injectability (bottom). **B)** Drug-hydrogel absorption profile: ATCK > ATKR while ATLG absorbs Ator instantly. **C)** Drug-hydrogel release profile: ATCK < <u>ATKR < ATLG</u> at t < 24h and ATCK < <u>ATLG < ATKR</u> at t > 24h. At 7d, Ator was released from ATCK at 2.2 ± 0.7%, ATKR at 11.2 ± 4.3%, ATLG at 7.8 ± 1.1%.

After COMSOL modelling, the diffusion coefficients of Ator absorbed in ATKR and ATCK hydrogels were 1(10^-8^) and 2(10^-8^) 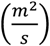, respectively (**Fig. 4A**). Their absorption concentration gradient was 147.2 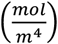. Multiplying each diffusion coefficient with the concentration gradients yields a diffusion flux of 1.47(10^-6^) and 2.94(10^-6^) 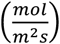 for ATKR and ATCK, respectively. They both have the same circular surface area as the Eppendorf flip-top tube (A = 4.52(10^-4^) m^2^, r = 12 mm). The cumulative mass absorbance rate is 6.66(10^-10^) 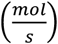 for ATKR and 13.32(10^-10^) 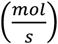 for ATCK. The diffusion coefficients of Ator released from ATKR, ATCK, and ATLG were 1.3(10^-12^), 1(10^-13^), and 1(10^-12^) 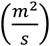, respectively (**Fig. 4B**). The drug release kinetics curve was not zero-order (linear), and further crosslinking agents can more effectively administer Ator.

**Fig 4.**
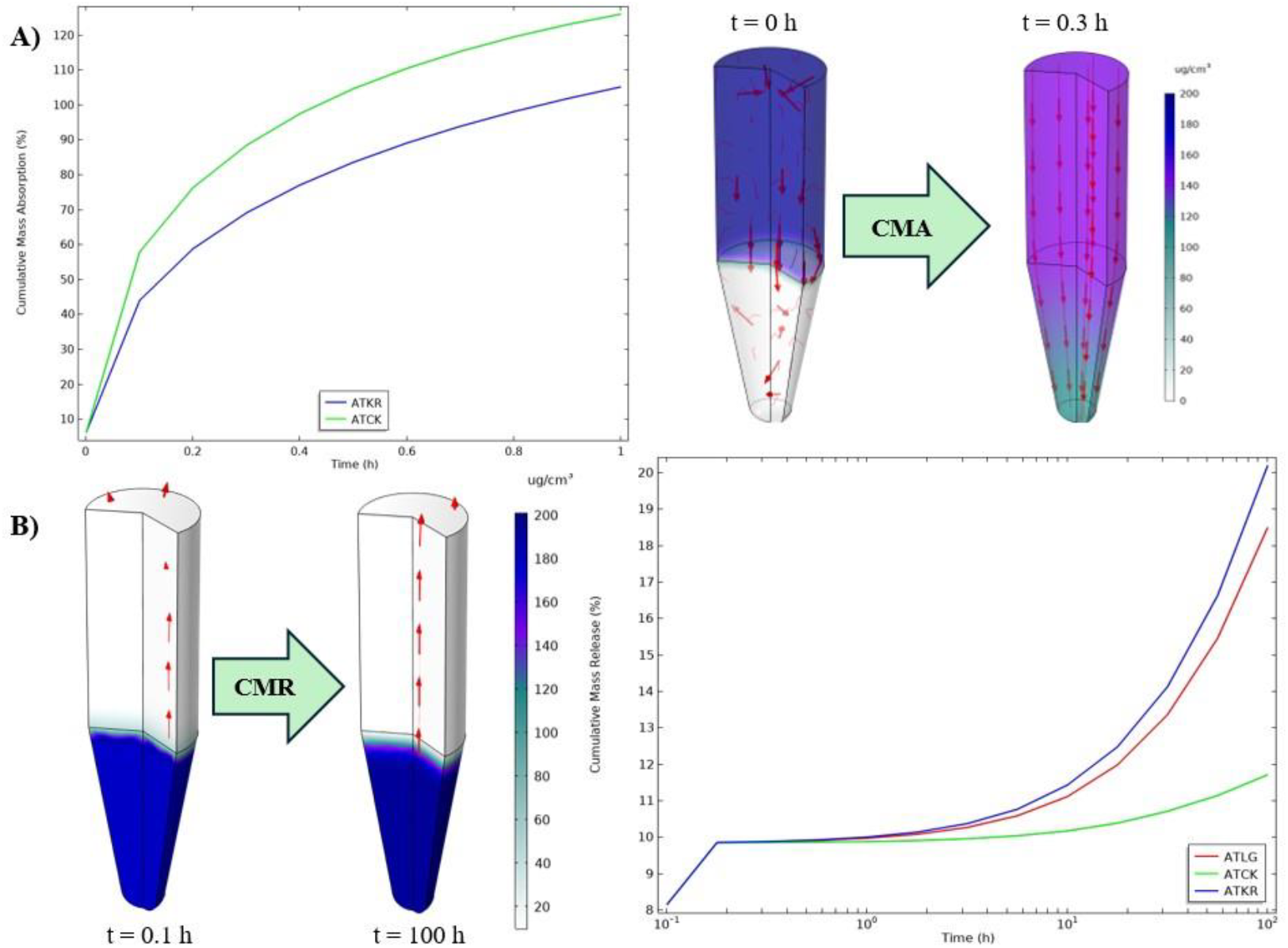
**A)** *Left*: Cumulative mass absorption curve of Ator in ATKR (blue) and ATCK (green). Ator immediately absorbs into calcium alginate spheres. *Right*: Model ATKR hydrogel before absorption (t = 0) and at 70% absorption (t = 0.3 h). **B)** *Left*: ATKR hydrogel releasing Ator back into PBS (t = 0.1 h) and slowly releases more Ator over time (m > 12% at t = 100 h).

The pharmacokinetic effects of Ator in a controlled environment was summarized in **Table 3** with the drug to hydrogel mass absorption was added to **Table 4**.

**Table 3.** Drug-hydrogel pharmacokinetics. ATKR = Ator in keratin gels. ATCK = Ator in keratin calcium gels. ATLG = Ator in calcium alginate gels. CMA = cumulative mass absorption. CMR = cumulative mass release. NA = Not applicable.

| Hydrogel<br>Groups | CMA<br>Diffusion<br>Coefficient<br>$\left(\frac{m^2}{s}\right)$ | CMR<br>Diffusion<br>Coefficient<br>$\left(\frac{m^2}{s}\right)$ | Diffusion<br>Flux $\left(\frac{mol}{m^2s}\right)$ | CMA Rate<br>$\left(\frac{mol}{s}\right)$ | CMR<br>(fold-<br>change) | |
| --- | --- | --- | --- | --- | --- | --- |
|  |  |  |  |  | 24 h | 7d |
| ATKR | 1*10<sup>-8</sup> | 1.3*10<sup>-12</sup> | 1.47*10<sup>-6</sup> | 6.66*10<sup>-10</sup> | 5 | 5 |
| ATCK | 2*10<sup>-8</sup> | 1*10<sup>-13</sup> | 2.94*10<sup>-6</sup> | 13.32*10<sup>-10</sup> | 1 | 1 |
| ATLG | NA | 1*10<sup>-12</sup> | NA | NA | 6.25 | 3.8 |
| Ator with no<br>gels | 1*10<sup>-6</sup> | 1*10<sup>-6</sup> | NA | NA | NA | NA |

**Table 4.** Biocompatibility and inhibition of drug-hydrogel interactions with MSC and L-929 cell lines. NA = Not applicable

| Hydrogel<br>Groups | Drug-Hydrogel Mass<br>Absorption (µg/mg) | Cell<br>Viability | Ator Effect (Fold-<br>change relative to<br>medium) | EC<sub>50</sub> (µM) |  |
| --- | --- | --- | --- | --- | --- |
| ATKR | 1.8 | < 70% | 106 ± 11% | NA |  |
| ATCK | 1.8 | 88.6 ± 34.7% | 228 ± 35% | NA |  |
| ATLG | 1.8 | < 70% | 124 ± 38% | NA |  |
| Ator with no<br>gels | NA | NA | NA | MSC | L-929 |
|  |  |  |  | 209 | 368 |

### 3.3 Cell Culture Biocompatibility and Inhibitory Responses

Testing hydrogel extracts for cell viability > 70% is the preferred ISO method, and after comparing positive and negative controls, ATCK proved to be non-cytotoxic with 88.6 ± 34.7% cell viability. The rest of the extracts had < 70% cell survival from this experiment, when similar previous cell culture experiments have shown high cell viability with our processed residual hair.

Low Ator doses did not affect the viability of MSCs, which enabled them to naturally proliferate and increase in number; but at relatively high increasing doses led to responsive inhibition (**Fig. 5, top**), with an EC_50_ of 209 μM (at similar range with fibro- and myofibroblasts, but with even higher EC_50_ using non-fibroblastic suggesting cell-specific actions). The same effects were observed with L-929 fibroblasts (**Fig. 5, bottom**), which exhibited responsive inhibition with an EC_50_ of 368 μM. Ator alone inhibits MSCs in cell culture better than fibroblasts, but the slow-release ATCK and ATLG do not affect these cells, and their proliferation is no different when compared to the control. Further studies can show that Ator conjugated in hydrogels can inhibit these cells. All gel groups inhibited the fibroblast cells similarly with no significant differences between gels with and without Ator. Further experimentation must be conducted to observe the effects of Ator on different fibroblasts that activate more easily since the quiescent FDA-approved ISO 10993-5 fibroblasts appear as better models for cytotoxicity tests than proliferation or inhibition experiments.

**Fig 5.**
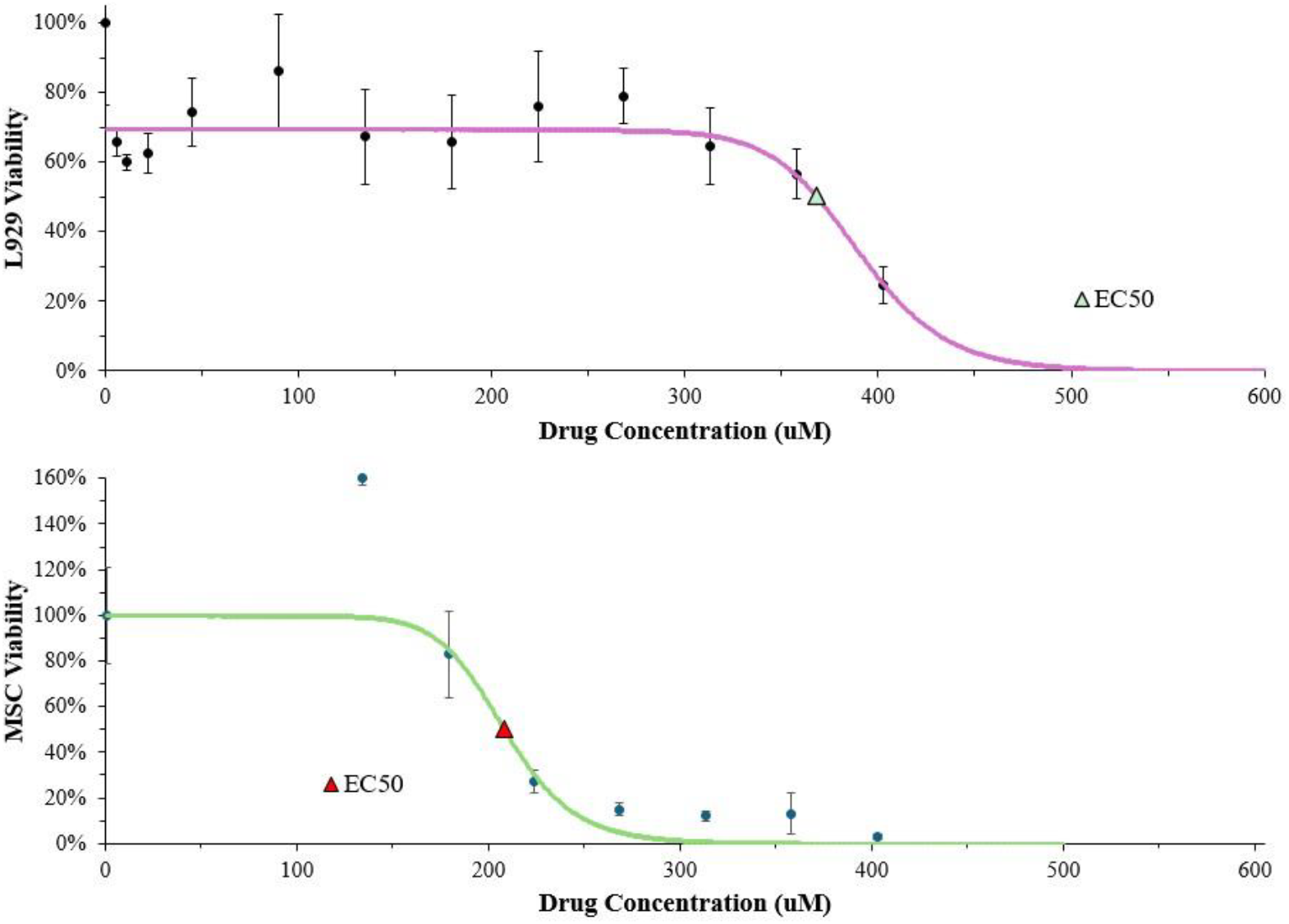
Negative sigmoidal inhibition curves of Ator on L-929 (top) and MSC (bottom) cell lines using Hill’s model with a negative Hill’s coefficient for toxicity of L-929 (n = -17.06) and MSCs (n = -11.55). L-929 EC_50_ = 368 μM. MSC EC_50_ = 209 μM.

Cell interactions and Ator’s effect with and without gel extracts are compiled in **Table 4**.

### 3.4 Experimental Synopsis

Quantitative and qualitative analysis of physical and pharmacological properties of keratin hydrogels and their extract cell viability are summarized in the following tables. Further biocompatibility tests are necessary to confirm the safety of cell-hydrogel interactions.

## 4. Discussion

### 4.1 Key Findings

After processing and reducing human hair meshes to shear the cuticle and expose the cortical contents, insoluble hair biomaterials hydrated into KTN hydrogels have desirable drug delivering properties such as integrated water content, high surface area to volume ratio, biodegradation, biomimicry and biocompatibility. Extracts with calcium and Ator were deemed safe and non-cytotoxic. Other hair biomaterials require further investigation and experimentation to retrieve accurate results. Although KRT hydrogels have more available negative charges to freely bond to the Ator calcium ions than CKRT gels do, the opposite was observed. As Ator began to conjugate to the negatively-charge KTN matrix, the higher Ca content absorbed the Ator faster in the ATCK hydrogel groups than in the ATKR ones. Similarly due to the higher concentration of Ca ions, the Ator was released slower from the ATCK hydrogels than ATKR. After exposing fibroblast and mesenchymal stem cell cultures to increasing concentrations of Ator, it is evident that the drug exhibits inhibitory effects, and further *in vivo* analysis is necessary to confirm the anti-fibrotic effects from reduced foreign body response and thinner fibrous encapsulation.

### 4.2 Mechanisms of Action

The KTN hydrogel acts as a matrix to control the release of Ator when applied topically or implanted surgically. The ratio between the hydrogel pore size and Ator molecular size directly affects the diffusion flux and drug release rate. The gel swell ratio also plays a large role in Ator release and is influenced by changes in the ion concentration in the microenvironment. Once released, Ator can enhance wound healing by signaling cell migration, collagen product, angiogenesis, and tissue remodeling while causing keratin proteins to detach from the gel matrix that also enhance these effects.

### 4.3 Advantages and Limitations

Ator sequestered in KTN hydrogels are derived from natural, abundant, biodegradable materials and exhibit effective localized drug delivery. KTN hydrogels have the potential to sequester different therapeutic agents for delivery at the same time, meaning they can manage diseases while also promoting regenerative properties. Some limitations of these drug carriers are the inconsistencies in biodegradability and release control. More experimental data is needed to determine the optimal gel hydration, degree of crosslinking, and environmental conditions such as temperature or pH. If the hydrogel degrades too quickly, it could lead to premature release of atorvastatin, reducing its effectiveness. Conversely, if it degrades too slowly, the drug may not be released effectively. Due to the novel concept of Ator sequestration in KTN hydrogels, there is limited preclinical and clinical data to support the safety and efficacy outside the cell culture hood.

### 4.4 Clinical Relevance and Future Directions

With its biocompatible natural origins, keratin hydrogels pose minimal risk of adverse immune responses. Their properties make them desirable for post-surgical healing, chronic wounds like diabetic ulcers, pressure wounds, and burns. Ator may have potential benefits in cosmetic dermatology for improving skin health, particularly in anti-aging treatments, and could provide a new approach to skin rejuvenation. With its primary cholesterol biosynthesis inhibitory functions, Ator may help to regulate lipid levels, which can be beneficial to manage conditions such as seborrheic dermatitis or other skin disorders linked to lipid metabolism.

Extensive preclinical and clinical studies are needed to validate the safety and efficacy of Ator-loaded KTN hydrogels for different applications. This would include randomized controlled trials to determine the impact on wound healing, inflammation, and skin regeneration, as well as assessing the long-term safety of topical and surgical Ator delivery. As the understanding of individual skin responses grows, personalized medicine and immunoengineering approaches can be developed to enhance the localized delivery treatment without invoking a large foreign body response. We provided computational simulations, but more advanced modeling techniques could help predict how Ator behaves in keratin-based hydrogels and result in more functional formulations. Understanding the bioavailability, penetration, and uptake mechanisms of Ator in different conditions could further enhance the therapeutic effectiveness.

## 5. Conclusions

Ator displays properties of inhibiting *in vitro* L-929 fibroblast and mesenchymal stem cell proliferation as its concentration increases. KTN hydrogels show capable sequestration and slow in vitro delivery of functional Ator, but more methods of crosslinking are necessary to determine the best way to administer the drug for the most effective outcomes. A follow-up *in vivo* mouse subcutaneous implantation study will be further explored to observe Ator’s fibrous encapsulation and foreign body response from KTN hydrogels.

## 6. Acknowledgements

We would like to thank Fusion Beauty Salon, Stylush Salon and Laser, Fresh Cuts, and Hair Designers for sources of human hair samples, Jason Williams, PhD for SEM assistance, Michelle Paszek and Ryan Radicone for gelation assistance, and Dr. Olga A. Carroll, Ph.D. for the inspiration, motivation, knowledge, and support.

